# CRISPR-Based Transcriptional Activation Tool for Silent Genes in Filamentous Fungi

**DOI:** 10.1101/2020.10.13.338012

**Authors:** László Mózsik, Mirthe Hoekzema, Niels A.W. de Kok, Roel A.L. Bovenberg, Yvonne Nygård, Arnold J.M. Driessen

**Affiliations:** Molecular Microbiology, Groningen Biomolecular Sciences and Biotechnology Institute, University of Groningen, Nijenborgh 7, 9747 AG, Groningen, The Netherlands; DSM Biotechnology Center, Alexander Fleminglaan 1, 2613 AX, Delft, The Netherlands; Synthetic Biology and Cell Engineering, Groningen Biomolecular Sciences and Biotechnology Institute, University of Groningen, Nijenborgh 7, 9747 AG, Groningen, The Netherlands; Chalmers University of Technology, Department of Biology and Biological Engineering, Division of Industrial Biotechnology, Kemivägen 10, SE-412 96, Gothenburg, Sweden; Swedish University of Agricultural Sciences, Department of Forest Mycology and Plant Pathology, Division of Forest Pathology, Almas allé 5, SE-756 51 Uppsala, Sweden

**Author notes:** These authors contributed equally to this work (László Mózsik, Mirthe Hoekzema).

**Keywords:** CRISPRa, dCas9, filamentous fungi, *Penicillium rubens*, secondary metabolites, biosynthetic gene clusters

## Abstract

Filamentous fungi are historically known to be a rich reservoir of bioactive compounds that are applied in a myriad of fields ranging from crop protection to medicine. The surge of genomic data available shows that fungi remain an excellent source for new pharmaceuticals. However, most of the responsible biosynthetic gene clusters are transcriptionally silent under laboratory growth conditions. Therefore, generic strategies for activation of these clusters are required. Here, we present a genome-editing-free, transcriptional regulation tool for filamentous fungi, based on the CRISPR activation (CRISPRa) methodology. Herein, a nuclease-defective mutant of Cas9 (dCas9) was fused to a highly active tripartite activator VP64-p65-Rta (VPR) to allow for sgRNA directed targeted gene regulation. dCas9-VPR was introduced, together with an easy to use sgRNA “plug-and-play” module, into an AMA1-vector, which is compatible with several filamentous fungal species. To demonstrate its potential, this vector was used to transcriptionally activate a fluorescent reporter gene under the control of the *penDE* core promoter in *Penicillium rubens*. Subsequently, we activated the transcriptionally silent, native *P. rubens* macrophorin biosynthetic gene cluster by targeting dCas9-VPR to the promoter region of the transcription factor *macR*. This resulted in the production of antimicrobial macrophorins. This CRISPRa technology can be used for the rapid and convenient activation of silent fungal biosynthetic gene clusters, and thereby aid in the identification of novel compounds such as antimicrobials.

## Introduction

Fungi are amongst the most proliferous producers of secondary metabolites (SMs). These molecules, while not intrinsically required for survival, provide a biological advantage to their host^1^. Many fungal SMs are beneficial to humankind and have a wide range of applications in human and animal healthcare (e.g. as antibiotics or immunosuppressants), food, agricultural and industrial sectors^2,3^. On the other hand, SMs can be toxic and some SMs contribute to the pathogenicity of fungi while others contaminate food and crops^4^. Genes involved in secondary metabolism are often arranged in clusters, so-called biosynthetic gene clusters (BGCs), and these are typically regulated by pathway-specific transcription factors. As more fungal genomes, and bioinformatics tools and databases (e.g. fungal antiSMASH^5^, MIBiG^6^) have become available for the prediction, annotation and prioritization of fungal BGCs, it has become clear that filamentous fungi have an even larger biosynthetic potential than previously anticipated.

Most of the BGCs are transcriptionally silent under laboratory growth conditions, therefore products of these clusters remain elusive^7^. Various methodologies have been developed for the activation of silent BGCs, including manipulation of both BGC specific as well as global transcriptional regulators, promoter-exchange, and heterologous expression in suitable host systems [reviewed in^8^]. Marker-free genome editing remains challenging, and with only a limited number of fungal selection markers available, extensive genome manipulations is a laborious task.

The bacterial CRISPR/Cas systems have emerged as versatile biotechnological tools^9,10^, and next to genome editing it can provide a promising alternative approach for transcriptional activation in fungi. CRISPR/Cas systems consist of only two components; a Cas nuclease and a programmable guide RNA. In case of the popular Cas9 system from *Streptococcus pyogenes* the protein can be guided to a genomic locus in a sequence-specific manner, using a single guide RNA (sgRNA) which consist of a short targeting crRNA sequence and the scaffold tracrRNA sequence. Methods for Cas9-based genome editing have been established in various filamentous fungal species [Reviewed^11,12^], including the industrially relevant fungi *Penicillium rubens^13,14^* (formerly identified as *P. chrysogenum*^15^). Cas9 and sgRNA delivery strategies include vector-based expression or genomic integration of transcriptional units encoding Cas9 and sgRNA. Alternatively, only Cas9 is expressed and the sgRNA is provided by a transformation of *in vitro* transcribed RNA, or both Cas9 and sgRNA are provided as pre-assembled ribonucleoprotein complexes (RNPs). The CRISPR/Cas9 genome editing tools established in filamentous fungi can edit the genome at a single as well as at multiple locations, and have effectively been applied in industrial fungi to improve compound production^11,12^.

Beyond genome editing, CRISPR/Cas can be used as a platform for RNA guided DNA-protein interactions, and thereby deliver various effector domains to a specific genomic location. By introducing point mutations into the two nucleolytic domains, nuclease deficient versions of Cas9, called dead Cas9 (dCas9), were created^16^. Because dCas9 binds in a sequence-specific manner, but does not cleave DNA, it can be used for transcriptional regulation^17,18^, epigenome editing^19,20^, visualization of a specific genomic locus^21^ and base editing^22^ in various eukaryotic species. For CRISPR/Cas mediated transcriptional activation (CRISPRa) several activating effector domains have been fused to dCas9 [reviewed in^23,24^]. The often-used VPR system consists of a three-component complex, four copies of the herpes simplex VP16 transactivation domain, the transactivation domain of nuclear factor kappa B, and Epstein-Barr virus R transactivator, VP64-p65-Rta, respectively^16^. dCas-VPR fusions have been successfully employed for upregulation of reporter and/or endogenous genes in mammalian cells^16^, in diploid^25^ and polyploid^26^ *Saccharomyces cerevisiae, Yarrowia lipolytica*^27^, *Candida albicans*^28^, and most recently also in the filamentous fungus *Aspergillus nidulans*^29^.

Here we report on the implementation of a dCas9-VPR-based genome editing free system for transcriptional activation in the filamentous fungus *Penicillium rubens*. We successfully utilized the CRISPRa tool to activate the cryptic macrophorin BGC, resulting in production of compounds with antimicrobial activity.

## Results

### Construction of a fungal CRISPRa tool

CRISPR/Cas mediated gene expression activation (CRISPRa) requires a catalytically dead CRISPR-associated protein (dCas) fused to an activation domain, as well as a sgRNA to guide it to the desired loci. Here, the widely utilized fusion of dCas9 from *Streptococcus pyogenes* to the tripartite activator, VP64-p65-Rta (VPR)^16^ was selected for activation. For easy implementation of CRISPRa in a broad range of filamentous fungi, we constructed an AMA1-based vector for expression of the NLS tagged dSpCas9-VPR under the 40S ribosomal protein S8 promoter (*p40S*) (Fig. 1a). The AMA1 sequence -originally isolated from *A. nidulans*-allows for autonomous vector replication in several filamentous fungal species^30,31^, and is often employed for Cas9 and sgRNA expression in gene-editing approaches in these organisms^11,13,32,33^. The AMA1 vector was also used to supply the sgRNA, establishing CRISPRa after a single transformation. The sgRNA was expressed from the constitutive *gpdA* promoter and flanked by hammerhead (HH) and hepatitis delta virus (HDV) ribozymes to ensure defined ends for sgRNA processing and optimal functionality (Fig. 1a)^34^.

**Figure 1.**
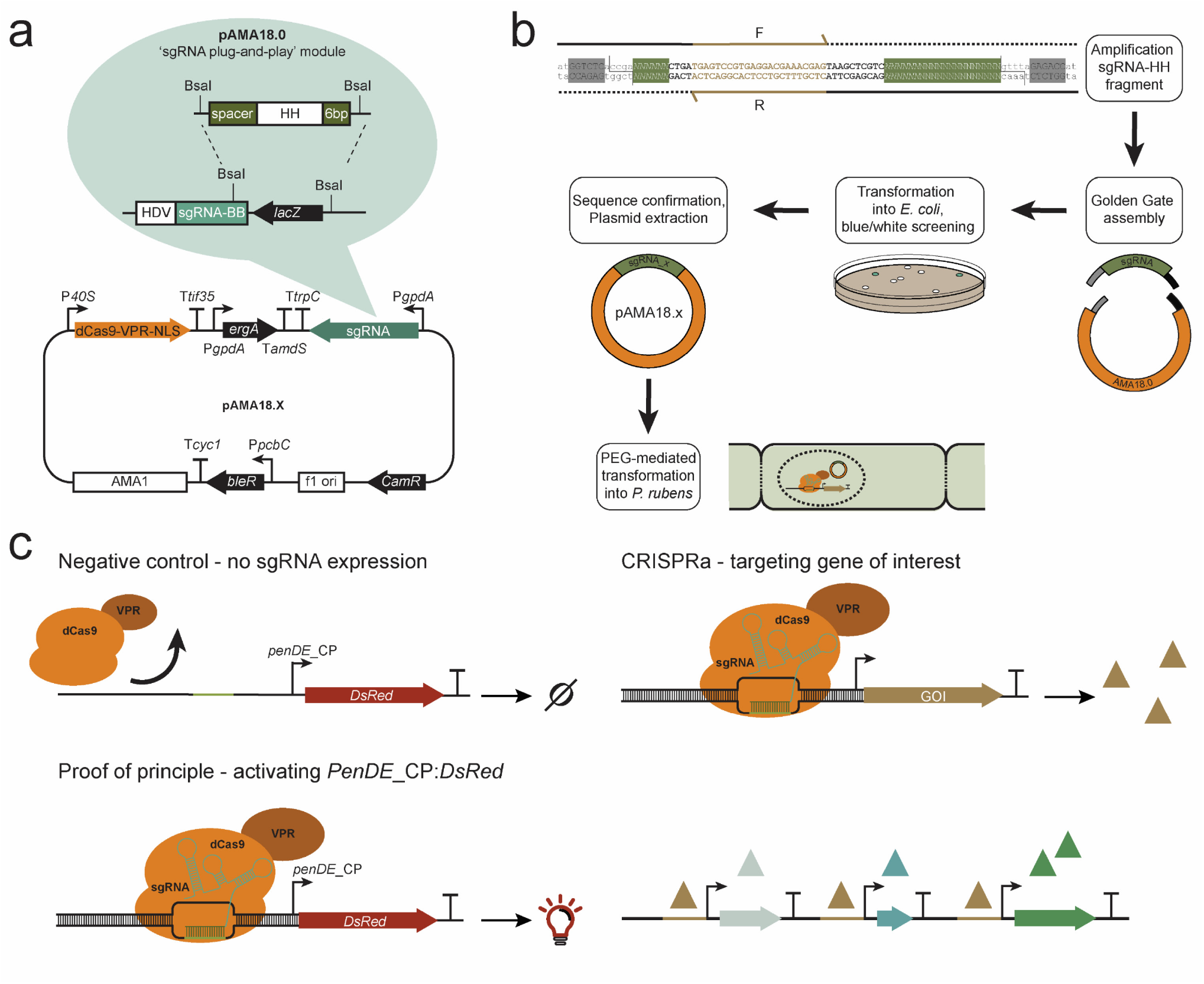
Overview of the programmable CRISPR/Cas-based transcriptional activation system implemented in *P. rubens*. (**a**) Schematic representation of the pAMA18.X-vector encoding the components of the CRISPR/Cas activation system, namely the dCas9-VPR and the ribozyme self-cleaved sgRNA. pAMA18.0 is the parent vector of all sgRNA containing vectors and contains the sgRNA “plug-and-play” module which is highlighted. (**b**) Diagram depicting the cloning strategy for insertion of the PCR amplified sgRNA into pAMA18.0. (**c**) CRISPRa proof of principle. In the control strain carrying pAMA18.0 no sgRNAs are transcribed, so while dCas9-VPR is present it is not targeted to specific loci and no transcriptional activation occurs. Correct targeting of the dCas9-VPR complex to the silent *penDE-CP* is leading to DsRed fluorescent protein expression and hence increased fluorescence. In the same fashion when dCas9-VPR is targeted to a promoter driving a gene of interest, this results in product formation. When the targeted promoter drives a transcriptional regulator this can result in activation/repression of multiple other genes, including entire BGCs.

Target specificity is determined by the sgRNA, thus by exchanging the sgRNA sequence different genes can be targeted for upregulation. To enable convenient and efficient exchange of sgRNA target sequences a sgRNA “plug-and-play” module was introduced into the AMA1 shuttle vector to facilitate cloning steps in *Escherichia coli* (Fig. 1a-b). This vector, which is the parent to all sgRNA expressing vectors, is called pLM-AMA18.0-dCas9-VPR (referred to as pAMA18.0) and also functions as a negative (non-targeting sgRNA) control. The sgRNA “plug-and-play” module works as follows; the chimeric sgRNA backbone sequence and the HDV ribozyme are already supplied on the AMA1-vector together with a *lacZ* gene flanked by BsaI restriction sites. The 20 nt spacer sequence defining the genomic target is supplied on a separate dsDNA molecule, together with the hammerhead ribozyme (HH) which includes the necessary 6 bp inverted repeat of the 5’-end of the spacer to complete the HH cleavage site. This dsDNA molecule can simply be created by PCR using two overlapping oligonucleotides (Fig. 1b) or alternatively ordered as chemically synthetized dsDNA. The fragment can then be inserted into pAMA18.0 using the Golden Gate cloning and the BsaI restriction sites^35^. As this removes the *lacZ* gene, positive bacterial clones can easily be detected with blue-white screening. After positive sequence verification and vector extraction, the created CRISPRa vector can be introduced into the filamentous fungi of choice (Fig. 1b).

### Proof of principle – activating penDE-CP_DsRed

In order to test if expression of dCas9-VPR and the sgRNA from the CRISPRa vector could activate transcription of a silent gene, we targeted dCas9-VPR to the *penDE* core promoter (*penDE*-CP). The 200 bp long *penDE-CP* has previously been shown to be functional, but insufficient to drive expression on its own^36^. For easy visualization of CRISPR based transcriptional activation, the *penDE-CP* was set to drive *DsRed-T1-SKL*, a red fluorescent reporter gene with peroxisomal targeting signal (Fig. 1c). The *penDE*-CP*_DsRed* reporter unit was integrated into the penicillin-locus of the *P. rubens* DS68530 (Δpenicillin-BGC), utilizing CRISPR/Cas9 ribonucleoprotein (RNP) facilitated co-transformation^13,14^ (Supplementary Fig. S1a).

Different pAMA18.0 derived vectors (pAMA18.a-f) expressing sgRNAs targeting loci +1 to −118 bp relative to the transcription start site (TSS) of the *penDE*-CP (Fig. 2a, Supplementary Fig. S2a, Supplementary Table S1) were transformed into *P. rubens* DS68530_*penDE*-CP*_DsRed* and strains were analyzed using fluorescence microscopy (Fig. 2b). Increased fluorescence intensity was seen in strains transcribing *penDE*-sgRNA_c, _d, _e and _f but not in strains transcribing *penDE*-sgRNA_a and _b. The DS68530_*penDE-CP_DsRed* strain carrying the pAMA18.0 negative control vector which did not express any sgRNA, showed only a minimal amount of fluorescence. DsRed expression was also evaluated using qPCR (Supplementary Fig. S3). These results confirm that activation of DsRed expression was CRISPRa dependent.

**Figure 2.**
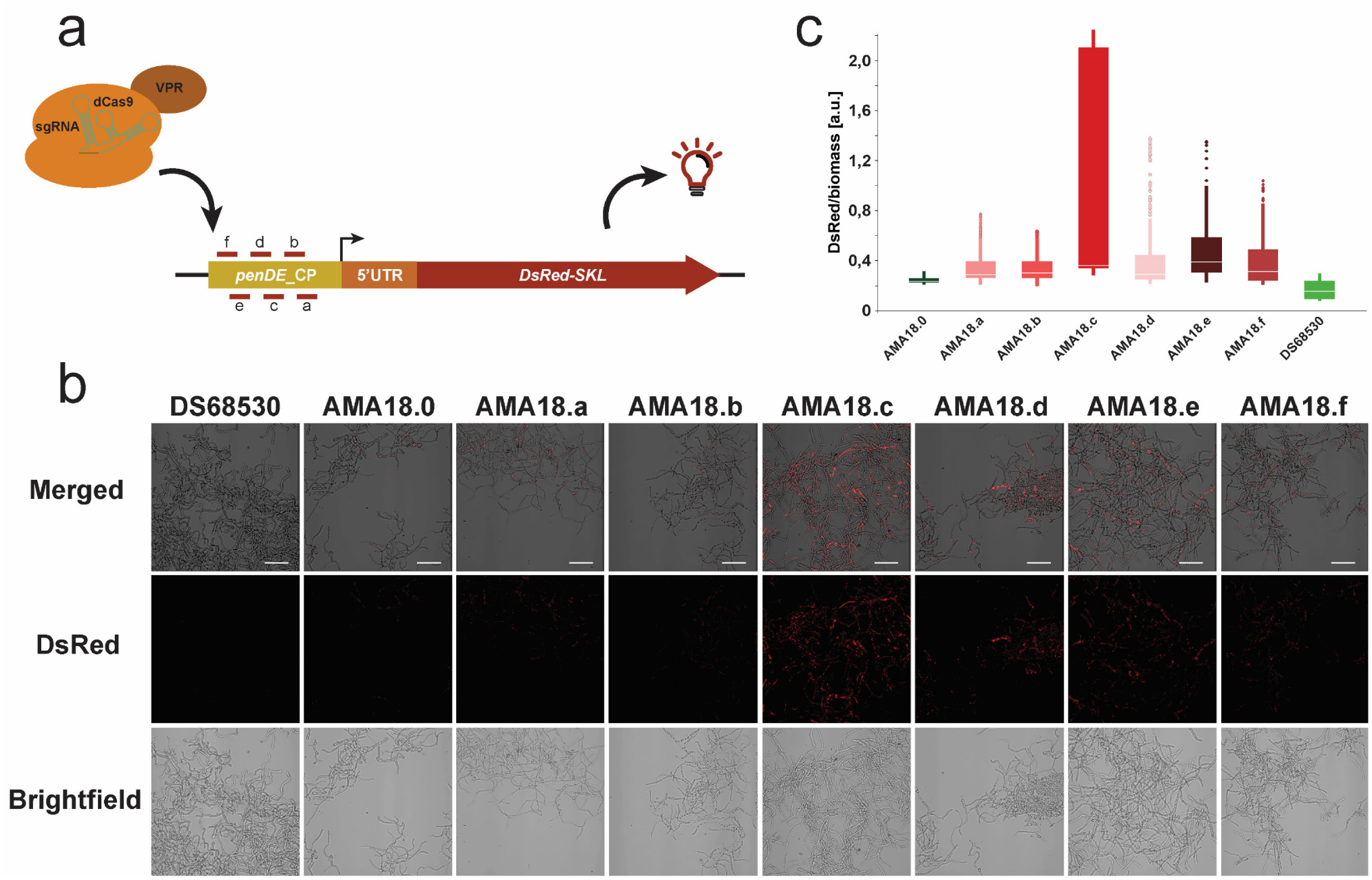
CRISPRa (dCas9-VPR) based activation of *penDE-CP_DsRed*. (**a**) Schematic representation of the *penDE-CP* upstream *DsRed*. The transcription start site (TSS) is indicated as a black arrow, short lines with letters indicate targeting sites of the sgRNAs. (**b**) Confocal fluorescence microscopy imaging of *DsRed* targeting CRISPRa strains and controls with no-sgRNA (AMA18.0) and without the *penDE-CP_DsRed* transcription unit (DS68530). Strains were grown for 5 days in liquid SMP media. Scale bars represent 50 μm. (**c**) Development of DsRed/biomass over time during time window of 0-40 hours cultivation in the BioLector system. Data were obtained from 3 separate experiments, each consisting of 2-3 biological replicates; error bars show the standard deviation.

To assess the performance of the different sgRNA target sequences, the BioLector microbioreactor system was used with online monitoring of scattered light (biomass) and red fluorescence intensity (Fig. 2c, Supplementary Fig. S4). The strength of DsRed activation by different sgRNAs was determined relative to biomass to avoid variance caused by small differences in growth. During the time interval of 0-40 h, DsRed/biomass values in CRISPRa strains were measured and compared to negative control strain carrying pAMA18.0 and the background fluorescence of the DS68530 parental strain. The pAMA18.c carrying strain showed the highest level of relative fluorescence and thus provided the most efficient activation compared to the non-sgRNA negative control (Fig. 2c). All other CRISPRa strains show activation of *penDE*-CP_*DsRed*, with weakest activation in strains carrying pAMA18.a and pAMA18.b vectors.

### A putative macrophorin gene cluster in P. rubens

Meroterpenoids represent a large family of natural compounds with diverse biological activities, such as the antimicrobial yanuthones found in *Aspergillus niger*^37,38^. Highly identical clusters have been found in *Penicillium* species^39^. These *Penicillium* BGCs contain an additional gene (*macJ*), which was shown in *Penicillium terreste* to encode a terpene cyclase responsible for cyclization of linear yanuthones leading to production of diverse macrophorin analogs^39^. The putative *P. rubens* macrophorin BGC consists of 11 biosynthetic genes, namely *macA-J* and *macR* as a transcriptional regulator of the cluster (Fig. 3a-b).

**Figure 3.**
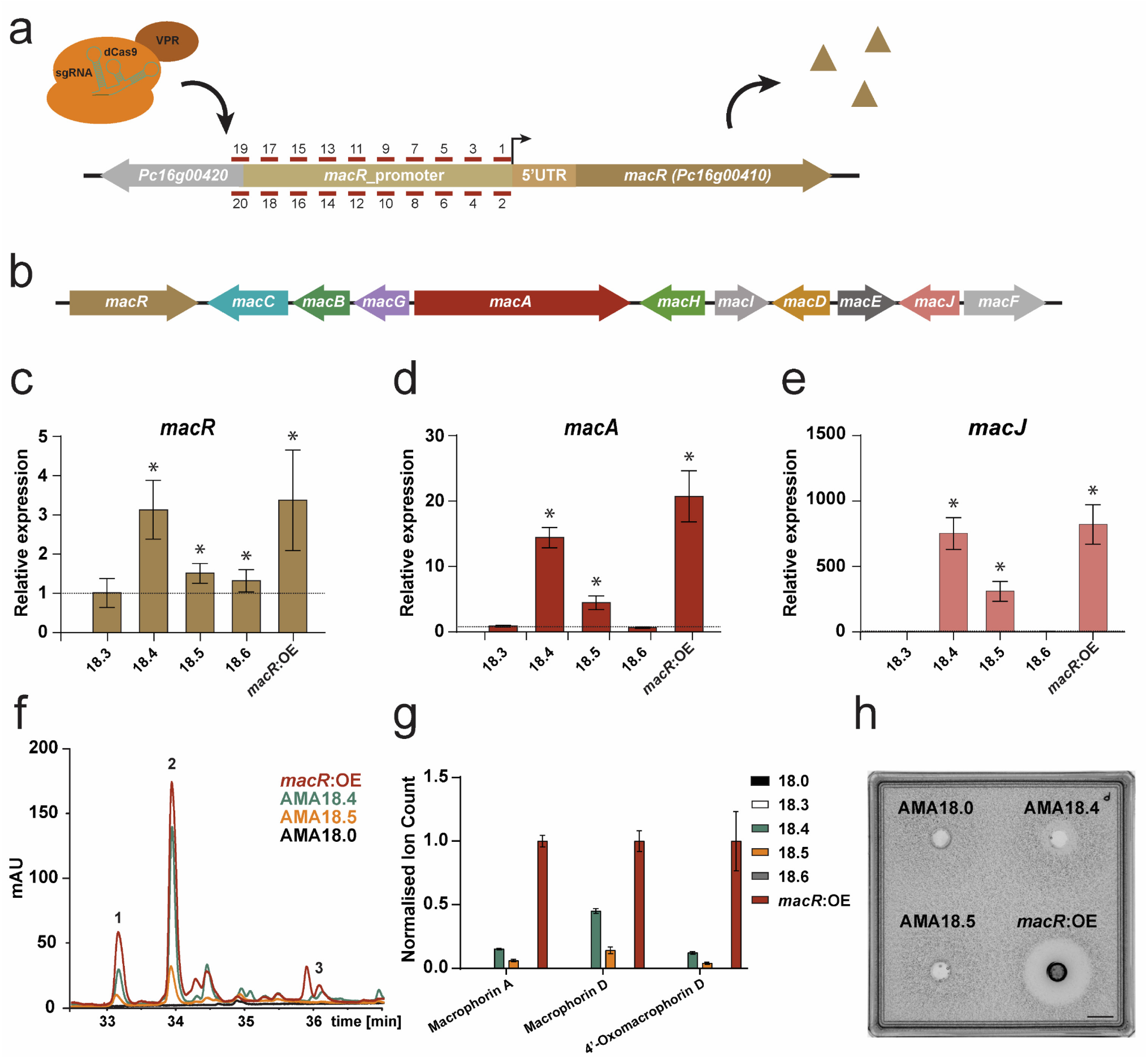
CRISPRa activation of the macrophorin BGC. **(a)** Schematic representation of the *macR* promoter region. The location of the putative transcription start site (TSS) is indicated as a black arrow, short lines with number indicate sgRNAs targeting sites. **(b)** Schematic representation of the Macrophorin BGC. qPCR analysis showing expression of *macR* **(c)** *macA* **(d)** and *macJ* **(e)** in the CRISPRa and *macR*:OE strains, relative to the strain carrying the pAMA18.0 vector (non-target control) after 5 days of growth in SMP medium. Error bars indicate the standard error of the mean of three biological replicates with two technical duplicates, and (*) indicates significant up-regulation (Student’s-test p-value <0.05). **(f)** LC-MS UV-VIS chromatogram (λ=700 nm) of hyphae extracts of CRISPRa and *macR*:OE strains representing macrophorin A (1), macrophorin D (2) and 4’-oxomacrophorin D (3). **(g)** LC-MS analysis of Macrophorin related compounds in hyphae extracts of CRISPRa and *macR*:OE strains. Error bars indicate the standard error of the mean of three biological replicates with two technical duplicates. **(h)** Bioassay to detect (macrophorin related) antimicrobial activity against *Micrococcus luteus* in the supernatant of indicated strains grown for 5 days in liquid SMP medium. The supernatant was concentrated 10-times and 100 μl was loaded in a well in top agar containing *M. luteus* at OD_600_ = 0.0125. Scale bar represents 10 mm.

Sequence alignment of the provisional sequence of *P. rubens macR* (*Pc16g00410*) to the *P. terrestre* LM2 *macR* coding sequence (*MF989995.1*) shows that the *P. rubens* sequence is predicted to have an additional intron leading to a premature stop codon. Without this intron, the *P. rubens macR* mRNA should produce a full-length product, similarly to *P. terrestre* LM2 *macR*. To test if *macR* codes for a functional protein we performed promoter replacement, substituting the promoter region of *macR* with the promoter of the *pcbC* (isopenicillin N synthase) gene (Supplementary Fig. S1b). The resulting increase in *macR* transcription (Fig. 3c) led to the activation of the cryptic BGC (Fig. 3d-e) and the production of macrophorins (Fig. 3f-g, Supplementary Table S2). We therefore conclude that *P. rubens macR* encodes for a functional transcription factor and that increased expression of *macR* leads to activation of the entire associated BGC without the need for additional transcription factors. Moreover, activation of this BGC leads to production of macrophorin-like compounds (Supplementary Table S2).

Sanger sequencing data of cDNA obtained from the *macR:OE* strain showed 2 introns in *P. rubens macR mRNA and* no pre-mature stop codon, in line with the coding sequence of *macR* of *P. terrestre* (*MF989995.1*) and the homologous *yanR* (*ASPNIDRAFT_44961*) of *A. niger*. It therefore seems likely *P. rubens* is capable of producing, not only functional, but also full length MacR. Additionally, a mutation (cDNA 2611C>T, P776S) mutation was identified in the ORF of *macR*. The effect of this mutation was not further investigated as *macR* remained capable of transcriptional activation. The sequence of *P. rubens* DS68530 *macR* cDNA can be found in Supplementary Figure S5.

### Activation of the transcriptionally silent macrophorin gene cluster

Since the *P. rubens* macrophorin BGC is silent under our growth conditions (Secondary Metabolite Producing [SMP] medium, 25 °C)^40,41^, it was selected for activation by CRISPRa. As no TSS is known for *macR*, 20 sgRNAs (MacR-sgRNA_1-20) were designed to target the entire 547 bp long, native promoter (Fig. 3a, Supplementary Table S1, Supplementary Fig. S2b). The *macR* targeting CRISPRa strains and the *macR:OE* positive control, were grown on SMP-agar for 10 days after which secondary metabolites were extracted from representative agar plugs, and analyzed by LC-MS (Supplementary Table S3). As expected, no macrophorin production was observed in the strain carrying the pAMA18.0 negative control with no sgRNA insert. Strains expressing MacR-sgRNA_4 and MacR-sgRNA_5 showed production of compounds with masses corresponding to macrophorin A (361.24 m/z [M+H]^+^), macrophorin D (505.28 m/z [M+H]^+^) and 4’-oxomacrophorin D (503.26 m/z [M+H]^+^) (Supplementary Table S2). None of the other CRISPRa strains showed exhibited macrophorin production.

Fungal strains carrying vector pAMA18.3-6 and pAMA18.0 were further investigated by qPCR (Fig. 3c-e) and metabolite profiling (Fig. 3f-g). As expected, strains carrying the pAMA18.4 or pAMA18.5 CRISPRa vector showed an increase in *macR* expression compared to the pAMA18.0 control, further confirming CRISPRa dependent transcriptional activation (Fig. 3c). The increase in *macR* expression resulted in transcriptional activation of the macrophorin BGC as exemplified by increased levels of *macA* (polyketide synthase) (Fig. 3d) and *macJ* (proposed terpene cyclase^39^) mRNA (Fig. 3e). In the strain carrying the pAMA18.4 vector, levels of transcriptional activation were comparable to those in the positive control *macR*:OE while strain carrying vector pAMA18.5 showed a ~3-fold lower transcription compared to this control, for all genes investigated (Fig. 3 c-e). No increased expression of *macR*, *macA* or *macJ* was observed for the strain carrying pAMA18.3. In the strain carrying vector pAMA18.6, a slight upregulation of *macR* was observed but this did not result in induction of *macA* and *macJ* (Fig. 3 c-e). In line with this, the strain carrying pAMA18.5 produced lower amounts of the examined macrophorin related metabolites compared to the strain with pAMA18.4 (Fig. 3 f-g). However, while qPCR analysis showed similar mRNA levels between the *macR*:OE and pAMA18.4 strains, compound production for macrophorin A and 4’-oxomacrophorin D was lower in AMA18.4 compared to the *macR*:OE strain. Strain AMA18.4 reached highest production for macrophorin D at ~38% of the ion intensity measured in *macR*:OE.

As the related yanuthones produced by *A. niger* display antimicrobial activity against gram positive bacteria^42^, we analyzed the activity of our macrophorin producing *Penicillium* strains against *Micrococcus luteus* using the agar diffusion method. The transformed parent strain *P. rubens* DS68530 does not contain the penicillin BGC, and consequently does not produce compounds inhibiting the growth of *M. luteus*. We observed a clearance zone around concentrated supernatant from the *macR*:OE strain grown for 5 days in SMP medium, and to a lesser extent also around that of the AMA18.4 strain, but not that of the control (AMA18.0) or the AMA18.5 strain (Fig. 3h). This indicates that the macrophorins produced by *P. rubens* are indeed bioactive against Gram-positive bacteria, and CRISPRa dependent activation of the BGC is sufficient to induce antimicrobial activity.

Interestingly, we observed a dark brownish pigmentation of the hyphae of the *macR:OE* strain after 5 days of cultivation on R-agar and SMP-agar as well as on day 1 in SMP liquid medium. The strain carrying the CRISPRa vector pAMA18.4 displayed a milder coloration compared to the colorless hyphae of the parent strain (Supplementary Fig. S6). Color formation in these *macR* over-expression strains was not investigated further.

## Discussion

In this work, we report the application of dCas9-VPR based CRISPRa in the ascomycetous filamentous fungus *Penicillium rubens*. While *Penicillium* is acclaimed for its production of β-lactam antibiotics, it harbors many more BGCs of which a substantial portion remain uncharacterized^43^. CRISPRa systems have been established in many model organisms as an ideal technology for transcriptional regulation and could aid in activating these often silent BGCs to facilitate characterization.

In our approach dCas9-VPR and the sgRNA are episomally encoded on the same AMA1-based vector, hence a single transformation is enough to establish CRISPRa in *Penicillium*. Moreover, because AMA1 supports autonomous vector replication in several filamentous fungal species^30,31^, and as we use established fungal promoters, terminators, and ribozyme based sgRNA processing, we expect the vector to be transferable to other fungal species. The sgRNA “plug-and-play” module of our CRISPRa vector combines Golden Gate cloning approach with blue/white screening. This allows for convenient cloning of new sgRNAs into the vector, reducing experimental time. This is especially important since general criteria for successful sgRNA design are difficult to define, and empiric testing of sgRNAs for each promoter region of interest remains necessary. Due to the ease of cloning our AMA1 vector, this CRISPRa technology could potentially be implemented in connection with larger scale fungal protoplast transformations using microtiter plates^44^, for example in combination with deploying multiple, separate sgRNA processing vectors in one transformation^45^.

To assess the CRISPRa vector for activation of transcriptionally silent promoter activation, we integrated a *penDE* core promoter driven *DsRed* gene into the penicillin-locus of *P. rubens* DS68530 (Δpenicillin-BGC). This *penDE-CP* was selected because it has been reported previously to be insufficient to express the fluorescent reporter on its own, instead depending on the presence of a synthetic transcription factor^36^. Fluorescence microscopy showed a clear increase in fluorescence with 4/6 sgRNAs tested, compared to a non-sgRNA expressing control (Fig. 2b). Quantification of fluorescence using a BioLector microbioreactor showed increased fluorescence for 6/6 sgRNA used, showing weak activation for *penDE*-sgRNA_a and *penDE*-sgRNA_b sgRNAs, and, in line with fluorescence microscopy results, *penDE*-sgRNA_c standing out as the most efficient activator (Fig. 2c). The discrepancy between fluorescence microscopy and the BioLector results could possibly be explained by a higher sensitivity of the BioLector, different cultivation method and time points (day 5 of shake-flask cultures for microscopy, average fluorescent during the first 40 hours for the BioLector cultivations).

In *A. niger*, Roux and co-workers observed that dCas9-VPR mediated activation of a mCherry fluorescent reporter fused to the transcriptionally silent *Parastagonospora nodorum elcA* promoter was stronger with sgRNAs targeting closer to the start codon, in a window of 162-342 bp upstream of the ATG ^29^. We target a region 106-170 bp upstream the start codon ATG (32-96 bp upstream the TSS) and observe the highest activation with *penDE*-sgRNA_c targeting 129 bp upstream the ATG, and the least with *penDE*-sgRNA_a and _b (not detectable by microcopy) targeting closer to the start codon. We thus do not see the same trend – stronger CRISPRa for sgRNAs targeting closer to the start codon – however we already target a window closer to the ATG compared to Roux *et al*.. This exemplifies that it remains difficult to define an optimal targeting conditions, and ideally several sgRNAs should be tested when establishing CRISPRa for a new promoter. In line with what previously was reported for *S. cerevisiae*, we did not observe an effect on CRISPRa efficiency when targeting the plus or minus strand ^46^.

To show our CRISPRa system can upregulate an entire silent BGC in *P. rubens* and induce metabolite production, we targeted the *macR* transcription factor of the endogenous macrophorin biosynthesis cluster. Macrophorins are a member of the meroterpenoids, a family of natural compounds which also include, for example, the antimicrobial yanuthones produced by *A. niger*^37,38^. Homologous macrophorin BGC have been identified in *Penicillium* species, and *P. terreste* has been shown to produce macrophorins, through the cyclization of yanuthones^39^. Out of the 20 sgRNAs tested, two resulted in transcriptional activation of the Macrophorin BGC (through the activation of transcriptional factor *macR*) (Fig. 2c-d) and secondary metabolite production (Fig. 2f-g). Although it is impossible to distinguish macrophorins and yanuthones with the method used as they have the same molecular formula, activation of the *macJ* terpene cyclase should lead to cyclic macrophorins^39^. Additionally, we could show that the supernatant of the CRISPRa activated strain grown five days in SMP media exhibited antimicrobial activity against the Gram-positive bacterium *M. luteus* (Fig. 3h). This clearly shows that our dCas9-VPR vector is capable of awakening silent BGCs in *Penicillium* and that the method can aid in product identification and characterization. It should be noted that exchanging the native *macR* promoter with the *pcbC* promoter resulted in higher compound production (Fig. 3g). It might therefore be beneficial to perform promoter exchange for high level production of interesting compounds identified using the CRISPRa technology. A possible explanation for why a larger proportion of the sgRNAs targeting *penDE-CP* (6/6) lead to transcriptional activation compared to *macR* (2/20) may be that the CP is free from most of its native regulatory elements, reducing chances of interference with the binding of the dCas9-VPR regulator. A limiting factor for this way of BGC activation is the need to identify a positive regulator for the cluster, which might not always be known. However, bioinformatics tools like antiSMASH^47^ or CASSIS^48^ could aid by identifying candidate regulators.

Recently, dCas12a (previously Cpf1), from *Lachnospiraceae bacterium* (dLbCas12a) or *Acidaminococcus* sp. (dAsCas12a), has become a popular alternative to dCas9 for gene regulation^49,50^. The Cas12a system has been popularized due to its ease of multiplexing; dCas12a uses smaller guide RNAs and is capable of processing these from a longer precursor CRISPR RNA^51^. Recent literature shows processing of 20 crRNA from a single precursor and simultaneous upregulation of 10 genes by dCas12a fused to a combination of the p65 activation domain together with the heat shock factor 1 in human embryonic kidney (HEK) 293T cells, exemplifying the potential of multiplex gene regulation using dCas12a^52^. A potential drawback for using dCas12a in fungi is the low activity at temperatures below 28 °C, while most fungal species grow optimally at temperatures between 25 °C and 30 °C. However Roux and co-workers recently engineered an temperature tolerant Cas12a mutant (dLbCas12a^D156R^-VPR), which was successfully employed for CRISPRa mediated gene activation in *A. nidulans* at 25 °C^29^. While dCas12a is an attractive choice when aiming to upregulate multiple genes simultaneously, for single target activation dCas9-VPR is still a good option. We got significant upregulation of an entire BGC using a single sgRNA targeting the TF of the BGC. For dLbCas12a based upregulation in *A. nidulans* (the unmutated dLbCas12a grown at 37 °C) multiple crRNAs were required for gene activation^29^. Another consideration when choosing a system is the different PAM requirement, NGG for (d)Cas9 and TTTN for (d)Cas12a. Depending on PAM availability in the genome one or the other could be preferable.

In conclusion we demonstrated that CRISPRa, specifically AMA1 vector-based expression of a dCas9-VPR fusion, can be used for the transcriptional activation of silent BGCs in *P. rubens*. We anticipate that the CRISPRa tool presented here can be widely used to awaken cryptic BGC in filamentous fungal species and thereby aid in the discovery of novel bioactive secondary metabolites.

## Methods

### Chemicals, reagents and oligodeoxyribonucleotides

All medium components and chemicals were purchased from Sigma-Aldrich (Zwijndrecht, Netherlands) or Merck (Darmstadt, Germany). Oligodeoxyribonucleotide primers (Supplementary Table S4) were obtained from Merck. Enzymes were obtained from Thermo Fisher Scientific (Waltham, MA) unless otherwise stated. For design of nucleic acid constructs, *in-silico* restriction cloning, Gibson Assembly and inspection of Sanger sequencing results, SnapGene (GSL Biotech) was used.

### Vector construction

The Golden Gate technology based Modular Cloning (MoClo) system^53^ using Type IIS BpiI and BsaI restriction enzymes were employed for the construction of all vectors unless stated otherwise. Constructed vectors with their destination vectors and corresponding PCR fragments or DNA donor vectors can be found in Supplementary Table S5. PCR amplifications were conducted using KAPA HiFi HotStart ReadyMix (Roche Diagnostic, Switzerland) according to the instructions of the manufacturer. Internal BsaI, BpiI recognition sites were removed for MoClo compatibility.

dCas9-2xNLS-VPR was amplified from two different sources. NLS-VPR was amplified from pAG414GPD-dCas9-VPR template (AddGene ID # 63801)^16^ and dCas9-NLS was amplified with adding D10A, D839A, H840A and N863A modifications from synthetic spCas9 pYTK036 template, provided as part of the Yeast MoClo Toolkit (AddGene ID # 65143)^54^.

The HH and HDV ribozyme based “Plug-and-Play” sgRNA transcription unit was amplified in three parts, where the *gpdA* promoter and the *trpC* terminator together with the HDV self-cleaving sequence were amplified from vector pFC334 (AddGene ID # 87846)^33^ and LacZ alpha gene was amplified from the MoClo ToolKit vector pICH41308 (AddGene ID # 47998)^55^. The promoter of 40S ribosomal protein S8 of *A. nidulans* (*AN0465.2*, referred to as *40S*) and the *tif35* terminator of *P. rubens Pc22g19890*, as well as the transcription unit *penDE-CP-DsRed-SKL*-T*Act*, were amplified from pVE2_10 (AddGene ID #154228)^36^. The terbinafine selection marker as Pgpda-ergA-TamdS transcription unit was amplified from pCP1_45^41^. The promoter of *pcbC* (*Pc21g21380*, IPNS) was amplified from pVE2_19 (AddGene ID #154241)^36^, adding 80 bp flanking regions for homologous recombination. The phleomycin selection marker was amplified from pDSM-JAK-109^56^ providing the P*pcbC-ble*-T*CYC1* transcription unit, adding 80 bp flanking regions for homologous recombination (Supplementary Fig. S1b, Supplementary Table S6).

Our autonomously replicating shuttle vector, carrying the AMA1 sequence, was based on pDSM-JAK-109^56^ where the *Pgpda-DsRed-SKL-TpenDE* transcriptional unit was removed using the BspTI and NotI restriction enzymes. The linear vector was treated with the Klenow Fragment and ligated to the circular vector using the T4 DNA Ligase according to the instructions of the manufacturer, creating a new AMA1 vector without *DsRed* expression. In order to create the CRISPRa vector, this vector was linearized with BspTI and was assembled by Gibson Cloning using PCR fragments G1, G2 G3 (Supplementary Table S5) carrying a terbinafine selection marker, dCas9-VPR and the sgRNA transcription unit respectively. CRISPRa vector pLM-AMA18.0 is deposited to AddGene under ID #138945. Parallel with this work a catalytically active spCas9 expressing vector was also established (pLM-AMA15.0 AddGene ID #138944) and utilized for genome editing [manuscript in preparation].

### sgRNA target design and cloning

Promoter sequences were analyzed with CCTop^57^ for possible CRISPR RNA guides with the following limitations: protospacer adjacent motif (PAM): NGG, target sequence length 20bp, core length 12bp, mismatches taken into account for prediction in core sequence 2, number of total mismatches 4 and using *P. rubens Wisconsin 54-1255* as the reference genome. Predicted protospacers were manually curated for minimizing off-target effects and selecting high CRISPRater^58^ scores.

Primers were designed to create 89 bp long dsDNA inserts by PCR, containing the unique 20 nt spacer sequence, the hammerhead ribozyme, the 6 bp inverted repeat of the 5’-end of the spacer sequence and the BsaI type II restriction enzyme recognition sites.

For cloning the inserts into the vector pAMA18.0 a modified MoClo protocol^53^ was used, using FastDigest BsaI (Thermo Fisher Scientific, Waltham, MA) restriction enzyme with an initial 10 min digestion, 50 cycles of digestion and ligation (37 °C for 2 min, 16 °C for 5 min), followed by a final digestion step and a heat inactivation step. Correctly assembled vectors were identified with blue-white screening and confirmed by sequencing. After positive sequence verification and vector extraction, the created pAMA18.X (where X stands for the sgRNA ID) CRISPRa vector was introduced into the fungal strain of choice (DS68530_*penDE*-CP_*DsRed* or DS68530) creating the CRISPRa fungal strain AMA18.X (Fig. 1b, Supplementary Table S6).

### Fungal strains and transformation

*P. rubens* strain DS68530^40^ (Δpenicillin-BGC, *ΔhdfA*, derived from DS17690) was kindly provided by Centrient Pharmaceuticals (former DSM Sinochem Pharmaceuticals, Netherlands). Protoplasts of *P. rubens* were obtained 48 hours post spore seeding in YGG medium and transformed using the methods and media described previously^14^.

### Media and culture conditions

Solid transformation medium was prepared using SAG solid medium (Sucrose 375 g/l: Agar 15 g/l; Glucose Monohydrate 10 g/l) supplemented, in this order, with 4 ml/l Trace Element Solution^59^, 25.7 ml/l stock solution A; 25.7 ml/l stock solution B and 2.4 ml/l 4 M KOH (where stock solution A contained the following: KCl 28.80 g/l; KH_2_PO_4_ 60.8 g/l; NaNO_3_ 240 g/l, at pH 5.5 (adjusted using KOH) and stock solution B contained: MgSO4·7H2O at 20.80 g/l). Selection for the terbinafine marker based *macR*:OE cassette and all CRISPRa vector carrying transformants was carried out using 1.1 μg/ml terbinafine hydrochloride (Sigma Aldrich) in the solid transformation medium. Terbinafine was supplemented in all media of consecutive experiments, whereas selection for *penDE*-CP_*DsRed* and P*pcbC*-*ble*-t*CYC*1 co-transformation was done using medium containing 50 μg/ml phleomycin (Invivogen, San Diego, CA). When the CRISPRa vector was transformed using PEG-mediated transformation, the amount of total DNA did not exceed 1 μg and no aurintricarboxylic acid was used. For each strain, 2 separate transformant colonies were selected as replicates and re-streaked individually on solid R-agar (see below) medium and cultivated for 7 days on 25 °C to produce spores, which were immobilized on lyophilized rice grains and used for further experiments. Schematic representation of engineering DS68530*_penDE-CP_DsRed* and *macR*:OE control strains, using CRISPR/Cas9 mediated homologous recombination based co-transformation into DS68530, is shown on Supplementary Figure S1. For each created strain, transformed DNA is listed in Supplementary Table S6.

For shake-flask liquid cultures, spores immobilized on lyophilized rice grains (0.2×10^6^ - 2×10^6^ spores/ml) were precultured for 24 hours in YGG medium^60^ before inoculation (1:7.5) into 30 ml Secondary Metabolite Producing (SMP) medium^60^ (Penicillin Producing Medium - PPM-without supplemented phenoxyacetic acid or phenylacetic acid), supplemented with 1.1 μg/ml terbinafine. Cultures were grown at 25 °C in a rotary incubator at 200 RPM for 5 days, after which mycelium was collected for total RNA extraction as well as extraction of secondary metabolites by vacuum filtration over filter paper. Solid R-agar medium^59^ was used for sporulation, SMP-agar (SMP medium supplemented with 15 g/l agar-agar) was used for cultivation, and for secondary metabolite extraction. All solid agar cultures were incubated at 25 °C.

### Total RNA extraction and cDNA synthesis

Total RNA was extracted from mycelium collected from cultures grown in SMP for 5 days at 25 °C. Wet biomass (~200 mg) was added to a screw cap tube containing 900 μl Trizol reagent (Thermo Fisher Scientific, Waltham, MA), 125 μl chloroform and glass beads (ø 0.75-1 mm, 500-600mg). The samples were stored at −80 °C until RNA isolation. The mycelium was disrupted using the FastPrep FP120 system (Qbiogene, Carlsbad, CA), followed by total RNA isolation using the phenol-chloroform extraction method. In short, after cell disruption phases were separated by centrifugation (10 min at 14000 x g, the upper phase was transferred to a new tube, followed by a chloroform extraction step (phase separation: 5 min at 12000 x g). RNA was precipitated by the addition of 1 volume isopropanol and incubated on ice for at least 10 minutes, followed by centrifugation (10 min at 12000 x g). Finally, the RNA was resuspended in milliQ H_2_O. DNAse treatment was done using the TURBO DNA-free Kit (Thermo Fisher Scientific, Waltham, MA), and the RNA concentration was determined using Nanodrop. cDNA was synthesized using RevertAid H Minus First Strand cDNA synthesis kit (Thermo Fisher Scientific, Waltham, MA) according to the manufacturer’s instructions for highly structured mRNAs using oligo (dT)_18_ primers and 1μg of total RNA as template in a 20 μl reaction.

### qPCR analysis

Primers used to analyze expression *DsRed, macR, macA* and *macJ* can be found in Supplementary Table S7. Primers were, when possible, designed to overlap an intron-exon junction to avoid amplification on gDNA. The *γ-actin* gene (*Pc20g11630*) was used as a control for normalization. The 25 μl qPCR reaction contained 4 μl of a 20x diluted cDNA synthesis reaction, 0.6 μM each of forward and reverse primer, and 12.5 μl SensiMix SYBR Hi-ROX master mix (Meridian Bioscience, Memphis, TN). Expression levels were determined with a MiniOpticon system (Bio-Rad, Hercules, CA) using the Bio-Rad CFX manager software, the threshold cycle (*Ct*) values were determined automatically by regression. Thermocycler conditions were as follows: 95 °C for 10 min, followed by 40 cycles of 95 °C for 15 sec, 60 °C for 30 sec, and 72 °C for 30 sec. Thereafter, a melting curve was generated to determine the specificity of the qPCRs.

### LC-MS sample preparation

For secondary metabolite analysis samples were taken from vacuum filtered mycelium or from solid SMP-agar medium as 3×1 cm diameter plugs. The plugs were transferred to a 4.0 ml glass vial and 1 ml acetone supplemented with 4 μl n-Dodecyl-β-D-maltoside (DDM) (10 mg/ml in methanol) was added as internal standard. The plugs were extracted ultrasonically for 60 min, after which the extracts were transferred to a clean vial and dried under a nitrogen stream at 25 °C. Dried extracts were resuspended in 200 μl methanol:milliQ-H2O (1:1) and filtered via a 0.2 μm PTFE syringe filter before the used for LC-MS analysis.

### LC-MS metabolite analysis

Metabolite analysis was performed using an Accella1250 UHPLC system coupled to a benchtop ESI-MS Orbitrap Exactive (Thermo Fisher Scientific, Waltham, MA) mass-spectrometer. A sample of 5 μl was injected onto a Waters Acquity CSH C18 UPLC (UHPLC) column (150×2.1 mm, 1.7 μm particle size) operating at 40 °C with a flow rate of 300 μl/min. Separation of the compounds was achieved by using a water-acetonitrile gradient system starting from 90% of solvent A (milliQ-water) and 5% solvent B (100% acetonitrile). 5% of solvent C (2% formic acid) was continuously added to maintain a final concentration of 0.1% of formic acid in the mobile phase. After 5 minutes of initial isocratic flow, the first linear gradient reached 60% of B at 30 minutes, and the second 95% of B at 35 minutes. A purge step for 10 minutes at 90% of B was followed by column equilibration for 15 minutes at the initial conditions. The column eluent was directed to a HESI-II ion source attached to the Exactive Orbitrap mass spectrometer operating at the scan range (m/z 80 – 1600 Da) and alternating between positive/negative polarity modes for each scan. LC-MS data were analyzed using the Thermo Scientific Xcalibur 2.2 processing software by applying the Genesis algorithm for peak detection and manual integration on the sum of the whole spectra of selected ions. The extracted ion counts of investigated compounds were normalized to the DDM internal standard and represented relative to the average detected values from the *MacR*:OE strain replicates. In addition to LC-MS only UV-VIS absorption was monitored at 220, 354 and 700 nm. Ions corresponding to the [M+H]^+^ pseudo molecular ions of the final steps of the macrophorin biosynthesis pathway (Macrophorin A, macrophorin D and 4’-oxomacrophorin D) were identified in chromatographic peaks (1), (2) and (3) respectively and were selected for further analysis. The peaks recorded by each channel for (1), (2) and (3) in match in retention time. The chromatogram recorded at 700 nm showed the best signal-to-noise ratio. (Fig. 3f, Supplementary Fig. S7). Due to the necessity of adding an in-line UV-VIS detector between the MS and the column to generate UV-VIS chromatograms, small discrepancy in Rt between different datasets was observed.

### Biolector

Spores (immobilized on 20 rice grains) were used to inoculate 10 ml SMP and cultures were incubated for 48 h in a rotary incubator at 200 rpm and 25 °C. For BioLector analysis and analysis of growth in FlowerPlate (MTP-48-B) wells, this pre-grown mycelium was diluted 8 times in fresh SMP, supplemented with 1.1 μg/ml terbinafine (except for parent strain DS68530). The 1 ml cultures were grown in the BioLector micro-bioreactor system (M2Plabs, Baesweiler, Germany), shaking at 800 rpm at 25 °C. In the BioLector, biomass was measured via scattered light at 620 nm excitation without an emission filter. The fluorescence of *DsRed-SKL* was measured every 30 min with “DsRed I” 550 nm (bandpass: 10 nm) excitation filter and 580 nm (bandpass: 10 nm) emission filter. Data were obtained from 3 separate experiments, each consisting of 2-3 biological replicates. The data obtained from the BioLector experiments were analyzed and presented using the TIBCO Spotfire Software (TIBCO Software Inc., Palo Alto, CA).

### Fluorescence microscopy

For visualization of DsRed-SKL fluorescent protein, liquid shake-flask cultures were cultivated for 5 days in SMP, and mycelium was collected and re-suspended in phosphate-buffered saline (58 mM Na2HPO4; 17 mM NaH_2_PO_4_; 68 mM NaCl, pH 7.3). Confocal imaging was performed on a Carl Zeiss LSM800 confocal microscope using 20x objective and ZEN 2009 software (Carl Zeiss, Oberkochen, Germany). The DsRed signal was visualized by excitation with a 543 nm helium neon laser (Lasos Lasertechnik, Jena, Germany), and emission was detected using a 565 to 615 nm band-pass emission filter^61^.

### Bio-assay

Macrophorin producing strains were tested for antimicrobial activity against *Micrococcus luteus* as follows: Supernatant of *P. rubens* strains carrying either the pAMA18.0, pAMA18.4, pAMA18.5 vector and the *macR:OE* strain was collected after 5 days of growth in liquid SMP medium and concentrated 10x in an Eppendorf Concentrator Plus (30 °C, vacuum for aqua solutions setting). An overlay of soft LA-agar (1%) inoculated with *M. luteus* to an OD_600_ of 0.125 was poured on top of an agar (1%) bottom layer with Oxford Towers (8×10 mm) spaced out evenly. The Oxford Towers were removed aseptically and 100 μl of the 10x concentrated supernatant was loaded in the resulting wells as indicated. The plate was incubated at 30 °C for 24 hours before imaging. The experiment was performed in triplicate.

## Abbreviations

BGC: Biosynthetic gene cluster
CRISPRa: CRISPR activation
SM: Secondary metabolite
cDNA: complementary DNA
sgRNA: chimeric single guide RNA
RNPs: ribonucleoprotein complexes
NLS: nuclear localization signal
CP: Core promoter
TSS: Transcription start site

## Author contributions

L.M. and M.H. contributed equally to this work. LC-MS analysis was carried out by N.A.W.K and L.M. BioLector analysis was carried out by Y.N.. L.M. designed the experiments. L.M. and M.H. carried out all other experiments and wrote the manuscript with critical feedback and help from N.A.W.K., R.A.L.B., Y.N., and A.J.M.D.. Y.N., R.A.L.B., and A.J.M.D. conceived the original idea.

## Acknowledgements

The authors would like to thank Arjen M. Krikken for the assistance with confocal fluorescence microscopy imaging.

## Additional information

Supplementary information is available for this paper at https://doi.org/xxx

