## Supplementary information for "CRISPR-Based Transcriptional Activation Tool for Silent Genes in Filamentous Fungi"

Table S1: CRISPR and CRISPRa sgRNA targeting sequences used in this study.

Table S2: Macrophorin related masses identified by LC-MS in *macR*:OE.

Table S3: LC-MS metabolite analysis of macrophorin related metabolites from agar plug extracts.

Table S4: Oligonucleotide primers used for strain and vector construction.

Table S5: Modular Cloning and Gibson Assembly based vector construction.

Table S6: Fungal strains created in this study.

Table S7: Oligonucleotide primers used for qPCR analysis.

Figure S1: CRISPR/Cas9 mediated engineering of *penDE*-CP_*DsRed* and *macR*:OE fungal strains.

Figure S2: Representation of sgRNAs targeting promoter sequences of *penDE*-CP and *macR*.

Figure S3: qPCR analysis on *DsRed* in CRISPRa strains.

Figure S4: Development of biomass, DsRed fluorescence and DsRed fluorescence in BioLector microbioreactors.

Figure S5: Complementary DNA sequence of *macR* (*Pc16g00410*) of *Penicillium rubens* DS68530.

Figure S6: Dark pigmentation of *Penicillium rubens* DS68530 hyphae in *macR* overexpression strains.

Figure S7: LC-MS UV-VIS chromatograms of hyphae extracts of *macR* overexpression strains.

**Table S1:** CRISPR and CRISPRa sgRNA targeting sequences with corresponding CRISPRater scores.

| Name | Sequence (5’-3’) | CRISPRater score (CCtop) |
|---|---|---|
| *PenDE*-sgRNA_a | GACCATAGCATGACACTGAT | 0.54 |
| *PenDE*-sgRNA_b | GTCCCATCAGTGTCATGCTA | 0.60 |
| *PenDE*-sgRNA_c | ATTGGCCGTAGCCACCAATC | 0.76 |
| *PenDE*-sgRNA_d | ATGCTATGGTCCCAGATTGG | 0.81 |
| *PenDE*-sgRNA_e | CACTGATGGGACATCAACTG | 0.28 |
| *PenDE*-sgRNA_f | GGATGCAGCAGGGATACTCG | 0.40 |
| *MacR*-sgRNA_1 | GACCATAGCATGACACTGAT | 0.73 |
| *MacR*-sgRNA_2 | GTCCCATCAGTGTCATGCTA | 0.69 |
| *MacR*-sgRNA_3 | ATTGGCCGTAGCCACCAATC | 0.60 |
| *MacR*-sgRNA_4 | ATGCTATGGTCCCAGATTGG | 0.84 |
| *MacR*-sgRNA_5 | CACTGATGGGACATCAACTG | 0.73 |
| *MacR*-sgRNA_6 | GGATGCAGCAGGGATACTCG | 0.71 |
| *MacR*-sgRNA_7 | AAAAACCCACCTCCCCTCAG | 0.78 |
| *MacR*-sgRNA_8 | ACTCCTCCTCTGAGGGGAGG | 0.70 |
| *MacR*-sgRNA_9 | CTGCATGCGAACTCCCAATA | 0.52 |
| *MacR*-sgRNA_10 | GGAGTTCGCATGCAGAGAAG | 0.77 |
| *MacR*-sgRNA_11 | CTCGCCCGGGAGGTAATTGG | 0.84 |
| *MacR*-sgRNA_12 | TTAGCCCCCAATTACCTCCC | 0.68 |
| *MacR*-sgRNA_13 | CAACTTCTCAAACATCGTTC | 0.66 |
| *MacR*-sgRNA_14 | GTTTGAGAAGTTGGTGTTGG | 0.67 |
| *MacR*-sgRNA_15 | GCTATAACACGAAGATGACA | 0.48 |
| *MacR*-sgRNA_16 | TAATAGGCAGGAAAGATTCG | 0.79 |
| *MacR*-sgRNA_17 | TATGTTTTGAAGTATACCCG | 0.73 |
| *MacR*-sgRNA_18 | ACATCTGTAGTGCTTACCTC | 0.69 |
| *MacR*-sgRNA_19 | GGATTTTTCACGATACGGGG | 0.76 |
| *MacR*-sgRNA_20 | CCCCGTATCGTGAAAAATCC | 0.61 |
| T7-pen-loci-editing[1] | GAACCAACATCATTAAGCAG | 0.70 |
| T7-*macR*:OE-editing[1] | AATGTTCCACTCCTCCTCTG | 0.85 |

**Table S2:** Macrophorin related masses identified by LC-MS in *macR*:OE from SMP-agar plug extracts after 10 days of growth.

| Nr | Compound | Formula | Theoretical mass m/z $[M+H]^+$ | Theoretical mass m/z $[M-H]^-$ | $t_{R*}$ (+ mode) | $t_{R*}$ (- mode) | Selected Rt for mass measurement | Detected mass m/z $[M+H]^+$ | Detected mass m/z $[M-H]^-$ | PPM error (+ mode) | PPM error (- mode) |
|---|---|---|---|---|---|---|---|---|---|---|---|
| 1 | Macrophorin A | $C_{22}H_{32}O4$ | 361.23734 | 359.22276 | 33.15 | 33.15 | 33.15 | 361.23776 | 359.22200 | **1.16** | **-2.12** |
| 2 | Macrophorin D | $C_{28}H_{40}O_8$ | 505.27959 | 503.26502 | 33.91 | 33.91 | 33.91 | 505.27999 | 503.26441 | **0.79** | **-1.21** |
| 3 | 4-Oxomacrophorin D | $C_{28}H_{38}O_8$ | 503.26394 | 501.24937 | 27.62/ 28.08/ 28.46/ 35.92 | 35.92 | 35.92 | 503.26469 | 501.24971 | **1.49** | **0.68** |
| 4 | DDM | $C_{24}H_{46}O_{11}$ | 511.31132 | 509.29672 | 28.12 | 28.12 | 28.12 | 511.3128 | 509.29678 | **2.89** | **0.12** |

**Table S3:** LC-MS metabolite analysis of macrophorin related metabolites from fungal SMP-agar plug extracts after 10 days of growth. Data represented as ion intensity normalized to added internal standard DDM.

| | Macrophorin A | Macrophorin D | 4'-Oxomacrophorin D |
|---|---|---|---|
| AMA18_0 | 0 | 0 | 1,436 |
| AMA18_1 | 0 | 0 | 12,470 |
| AMA18_2 | 0 | 0 | 4,473 |
| AMA18_3 | 0 | 0 | 5,843 |
| AMA18_4 | 117,925 | 117,925 | 80,451 |
| AMA18_5 | 771,392 | 77,170 | 52,530 |
| AMA18_6 | 2,646 | 0 | 1,989 |
| AMA18_7 | 3,456 | 0 | 418 |
| AMA18_8 | 0 | 827 | 1,157 |
| AMA18_9 | 1,391 | 449 | 0 |
| AMA18_10 | 0 | 0 | 3,899 |
| AMA18_11 | 800 | 0 | 1,947 |
| AMA18_12 | 0 | 0 | 2,857 |
| AMA18_13 | 0 | 0 | 5,682 |
| AMA18_14 | 0 | 636 | 1,071 |
| AMA18_15 | 0 | 811 | 3,323 |
| AMA18_16 | 0 | 0 | 5,503 |
| AMA18_17 | 0 | 764 | 1,491 |
| AMA18_18 | 956 | 0 | 806 |
| AMA18_19 | 0 | 1,321 | 0 |
| AMA18_20 | 0 | 1,691 | 3,113 |
| *MacR*:OE | 3,675,166 | 2,747,993 | 91,620 |

**Table S4:** Oligonucleotide primers used for strain and vector construction.

| Part ID | Description | Template | Primer Pair Sequences (5'->3') |
|---|---|---|---|
| A | *penDE*-CP_*DsRed-T1-SKL*-*Tact* (80-800 bp flanks) | pVE2_10[2] AddGene ID #154228 | F:TCGACACGCTTTACGAATTCCCATGG<br>R:GATATGCCGTCTGCAGAGACTGCGATA |
| B | P*pcbC-ble-*T*cyc1* (80bp flanks) | pJAK-109[3] | F:AGACTCGGTGATGCAGCAAATAGCGACTGTTCGTTGCGGGGTCCGAACCCGCTCGGCAGCACCGGG CTCTCCCTACTATCCCTCGA TAGCAGTCGACTACATGTATCTGCATGTTGCATC<br>R:TGACACTGATGGGACATCAACTGGGGCACCTCGAGTATCCCTGCTGCATCCGCCTAGTCTCTCCCCA TGGGAATTCGTAAAGCGTGTCGATAAGCTTGCAAATTAAAGCCTTCGAGCG |
| C | P*pcbC-macR* (80bp flanks) | pVE2_19[2] AddGene ID #154241 | F:GCTGCATTGGTCTGCCATTGC<br>R:CGAACCTTGCGTCGTCGGCAGACGATGCAACTGAGCGATAACCGGGGTTTCTTGATTTGGCGGGGC TCGGATAAAGGCATTGGTGTCTAGAAAAATAATGGTGAAAACTTG |
| D | *PgpdA-ergA*-T*amdS* terbinafine marker (80bp flanks) | pCP1_45[4] | F:CGCCCTGCTATCCCAACCCTGCACTTGTCCTCTTCTCTGCATGCGAACTCCCAATATGGTCACGATAA AAACCCACCTCCTACCGCTCGTACCATGGGTTGAG<br>R:CAGTGCTTCAGTCGCCCAGATTCTCGATGGAGATTGGCCAGGTCAGCCATATATACCCTGCAATGGC AGACCAATGCAGCGAATTCGAGCTCGGAGTGGATCC |
| E1 | Cas9m4-VPR-1 | pYTK036[5] AddGene ID #65143 | F:TGAAGACTTAATGGACAAGAAGTATTCTATCGGACTGGCCATCGGGACTAATAG<br>R:TGAAGACTTGCAGCCACGTCGTAGTCTGAGAGC |
| E2 | Cas9m4- VPR-2 | Annealed oligos | F:TGAAGACTTCTGCCATCGTCCCTCAGAGCTTCCTCAAAGACGACTCAATTGACAATAAGGTGCTGACT CGCTCAGACAAGGCCAAGTCTTCA<br>R:TGAAGACTTGGCCTTGTCTGAGCGAGTCAGCACCTTATTGTCAATTGAGTCGTCTTTGAGGAAGCTCT GAGGGACGATGGCAGAAGTCTTCA |
| E3 | Cas9m4- VPR-3 | pYTK036[5] AddGene ID #65143 | F:TGAAGACTTGGCCCGGGGAAAGTCAGATAACGTGC<br>R:TGAAGACTTATCCCCTCCGAGCTGTGAGAGG |
| E4 | Cas9m4- VPR-4 | pAG414GPD-dCas9-VPR[6] AddGene ID #63801 | F:GCTGAAGACTTGGATAGCAGGGCTGACCCCAAGAAGAA<br>R:TGAAGACTAAAGCTCAAAACAGAGATGTGTCGAAGATGGACAGT |
| F1 | HH-sgRNA-HDV “plug-and-play” 1 | pFC334[7] AddGene ID #87846 | F:CGGTCTCTAGCGGCGTAAGCTCCCTAATTGGCCC<br>R:CGGTCTCATCGGTGATGTCTGCTCAAGCGG |
| F2 | HH-sgRNA-HDV “plug-and-play” 2 | pICH41308[8] AddGene Kit ID #1000000044 | F:GAAGACTCCCGACGAGACCCAGCTGGCACGACAGGTTTC<br>R:GAAGACAAAAACGGAGACCACAGCTTGTCTGTAAGCGGATG |
| F3 | HH-sgRNA-HDV “plug-and-play” 3 | pFC334[7] AddGene ID #87846 | F:TGGTCTCAGTTTTAGAGCTAGAAATAGCAAGTTAAAATAAGGCTAG<br>R:CGGTCTCAGGAGGAGCCAAGAGCGGATTCCTCAGTCTCGTACGTCTC |
| G1 | Gibson unit 1, dCas9m4-VPR | pLM1_135 | F:GAATTCCTGCAGCCCCAGATCATCCTGTCTTCAGTCTTAACGCTGCAAGAATTCAAGCTTGGAG<br>R:TGGGATGTTCCATGGTAGCTGTGAA |
| G2 | Gibson unit 2, p*40S* flank with *PgpdA-ergA-TamdS* | pCP1_135 | F:CAAGGTTCTTCTCGAAGTAGTTGTTCT<br>R:CGCTCGTACCATGGGTTGAG |
| G3 | Gibson unit 3, sgRNA “plug-and-play” | pLM1_135 | F:ACAGGTGACTCTGGATGGC<br>R:ACCTTCAATATCAACTCTTTCAGGGGGGGAGCGGCCTTAAGTCGGCAACGAGAGGTATGTCTAAAGT |
| H | T7-sgRNA-transcription template[1] (penicillin-loci) | overlapping oligonucleotides | F:ATGTAATACGACTCACTATA**gAACCAACATCATTAAGCAG**GTTTCAGAGCTATGCTGGAAA*<br>R:AAAAAAGCACCGACTCGGTGCCACTTTTTCAAGTTGATAACGAACTAGTCTTATTTCAACTTGCTATG CTGTTTCCAGCATAGCTCTGAAAC |
| I | T7-sgRNA- transcription template[1] *macR*-OE | overlapping oligonucleotides | F:ATGTAATACGACTCACTATAg**AATGTTCCACTCCTCCTCTG**GTTTCAGAGCTATGCTGGAAA*<br>R:AAAAAAGCACCGACTCGGTGCCACTTTTTCAAGTTGATAACGAACTAGTCTTATTTCAACTTGCTATG CTGTTTCCAGCATAGCTCTGAAAC |

*20bp sgRNA target sequence shown in bold, lowercase “g” indicates T7 transcription site

**Table S5:** Modular Cloning and Gibson Assembly based vector construction.

| **Created vector** | **Description** | **Cloned Part IDs or MoClo units** | **Recipient vector** |
|---|---|---|---|
| LM0_36 | dCas9m4-2xNLS-VPR | E1, E2, E3, E4 | pICH41308[8] |
| LM1_100 | P*40S*-dCas9m4-2xNLS-VPR-T*tif35* | pZB0_21[2], pLM0_36, pYN0_10[2] | pICH47742[8] |
| LM1_113 | sgRNA “plug-and-play” transcription unit (P*gdpA*-lacZ-HDV-T*trpC*) | F1, F2, F3 | pICH47761[8] |
| LM2_135 | P*40S*-dCas9m4-2xNLS-VPR-T*tif35*, P*gpdA*-*ergA*, sgRNA “plug-and-play” transcription unit, PenFlanks, MoClo End-Linker on LVL2 MoClo backbone vector | pZB1_1[2], pLM1_100, pCP1_45[4], pLM1_113, pZB1_2[2], pICH41800[8] | pICH50505 (alternative of pAGM4673[8]) |
| pAMA18.0* | P*40S*-dCas9m4-2xNLS-VPR-T*tif35*, P*gpdA*-*ergA*-T*amdS*, sgRNA “plug-and-play” transcription unit on AMA1 backbone vector | G1, G2, G3 | pJAK-109 based linearized AMA1 vector[3] |

*Vector constructed using Gibson Assembly

**Table S6:** Fungal strains created in this study**.**

| Strain ID | Transformed DNA | Transformed strain | Transformation method |
|---|---|---|---|
| *penDE*-CP_*DsRed* | Part ID A and B | DS68530 | PEG mediated RNP-based CRISPR-Cas9 editing by homologous recombination[1,9] sgRNA transcribed using T7 polymerase from DNA of “Part ID: H” |
| *macR*:OE | Part ID C and D | DS68530 | PEG mediated RNP-based CRISPR-Cas9 editing by homologous recombination sgRNA transcribed using T7 polymerase from DNA of “Part ID: I” |
| AMA18.0_*DsRed* (no-sgRNA control for *DsRed*) | pAMA18.0 | DS68530_*penDE*-CP_*DsRed* | PEG mediated vector transformation |
| AMA18.a | pAMA18.a | DS68530_*penDE*-CP_*DsRed* | PEG mediated vector transformation |
| AMA18.b | pAMA18.b | DS68530_*penDE*-CP_*DsRed* | PEG mediated vector transformation |
| AMA18.c | pAMA18.c | DS68530_*penDE*-CP_*DsRed* | PEG mediated vector transformation |
| AMA18.d | pAMA18.d | DS68530_*penDE*-CP_*DsRed* | PEG mediated vector transformation |
| AMA18.e | pAMA18.e | DS68530_*penDE*-CP_*DsRed* | PEG mediated vector transformation |
| AMA18.f | pAMA18.f | DS68530_*penDE*-CP_*DsRed* | PEG mediated vector transformation |
| AMA18.0 (no-sgRNA control for *macR*) | pAMA18.0 | DS68530 | PEG mediated vector transformation |
| AMA18.1 | pAMA18.1 | DS68530 | PEG mediated vector transformation |
| AMA18.2 | pAMA18.2 | DS68530 | PEG mediated vector transformation |
| AMA18.3 | pAMA18.3 | DS68530 | PEG mediated vector transformation |
| AMA18.4 | pAMA18.4 | DS68530 | PEG mediated vector transformation |
| AMA18.5 | pAMA18.5 | DS68530 | PEG mediated vector transformation |
| AMA18.6 | pAMA18.6 | DS68530 | PEG mediated vector transformation |
| AMA18.7 | pAMA18.7 | DS68530 | PEG mediated vector transformation |
| AMA18.8 | pAMA18.8 | DS68530 | PEG mediated vector transformation |
| AMA18.9 | pAMA18.9 | DS68530 | PEG mediated vector transformation |
| AMA18.10 | pAMA18.10 | DS68530 | PEG mediated vector transformation |
| AMA18.11 | pAMA18.11 | DS68530 | PEG mediated vector transformation |
| AMA18.12 | pAMA18.12 | DS68530 | PEG mediated vector transformation |
| AMA18.13 | pAMA18.13 | DS68530 | PEG mediated vector transformation |
| AMA18.14 | pAMA18.14 | DS68530 | PEG mediated vector transformation |
| AMA18.15 | pAMA18.15 | DS68530 | PEG mediated vector transformation |
| AMA18.16 | pAMA18.16 | DS68530 | PEG mediated vector transformation |
| AMA18.17 | pAMA18.17 | DS68530 | PEG mediated vector transformation |
| AMA18.18 | pAMA18.18 | DS68530 | PEG mediated vector transformation |
| AMA18.19 | pAMA18.19 | DS68530 | PEG mediated vector transformation |
| AMA18.20 | pAMA18.20 | DS68530 | PEG mediated vector transformation |

**Table S7:** Oligonucleotide primers used for qPCR analysis.

| Gene | Sequences (5'->3') |
|---|---|
| *DsRed-T1-SKL* | F:CCAAGGTGTACGTGAAGCAC<br>R:CCTTGTAGATGAAGGAGCCGT |
| *macR* (*Pc16g00410*) | F:GACGACGCAAGGTTCGCT<br>R:GTCCTGCGGTGATACTGGTC |
| *macA* (*Pc16g00370*) | F:CGGGTTCAAACACGTCCGTA<br>R:CATCCAGTGCAACGCTAGGA |
| *macJ* (*Pc16g00320*) | F:TCTTGGGGAATTTGGTGGACA<br>R:CAGACCCGATACTACCCAGC |
| *γ-actin* (*Pc20g11630*) | F:CTGGCGGTATCCACGTCACC<br>R:AGGCCAGAATGGATCCACCG |

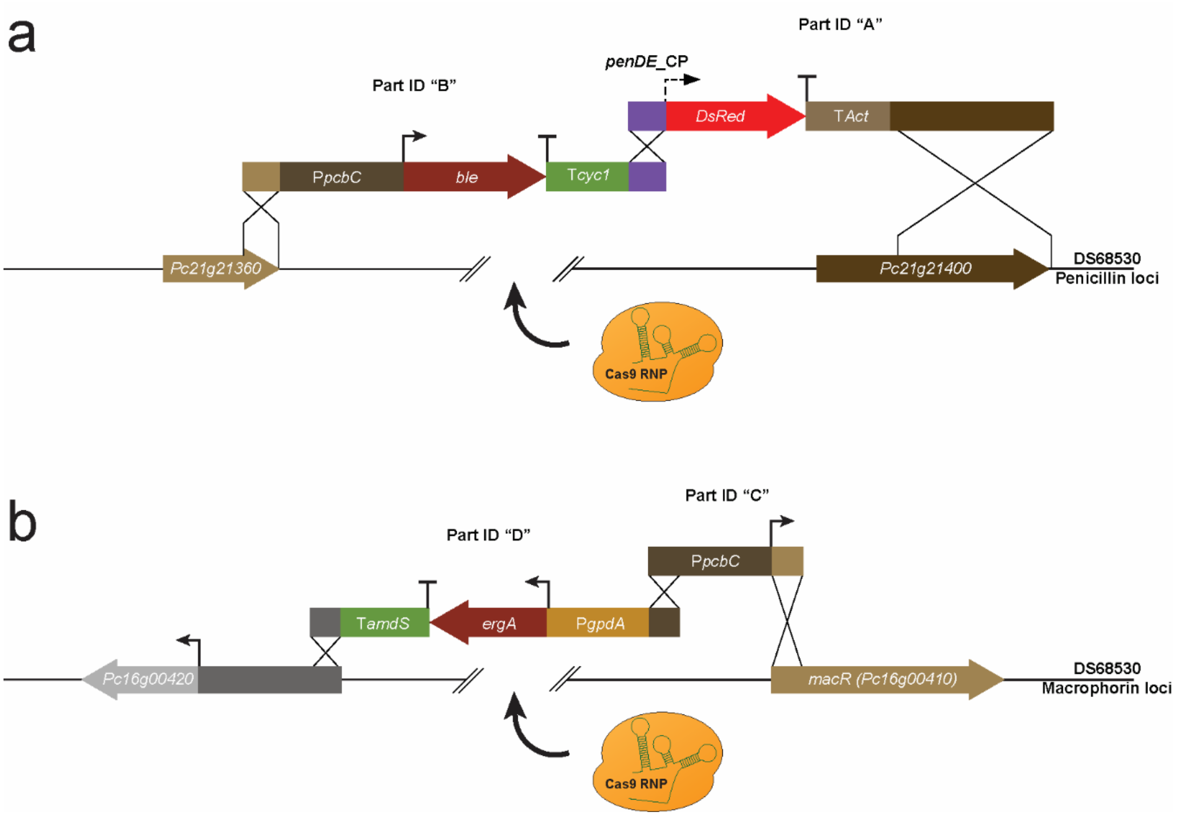


**Figure S1:** Schematic representation of CRISPR/Cas9 mediated co-transformation into DS68530 and engineering of *penDE*-CP_*DsRed* **(a)** and *macR*:OE **(b)** fungal strains, using Part ID A,B and C,D respectively. The *ergA* (terbinafine) or *ble* (phleomycin) marker provide selection and flanking regions for recombination with the marker-free cassette and the genomic DNA.

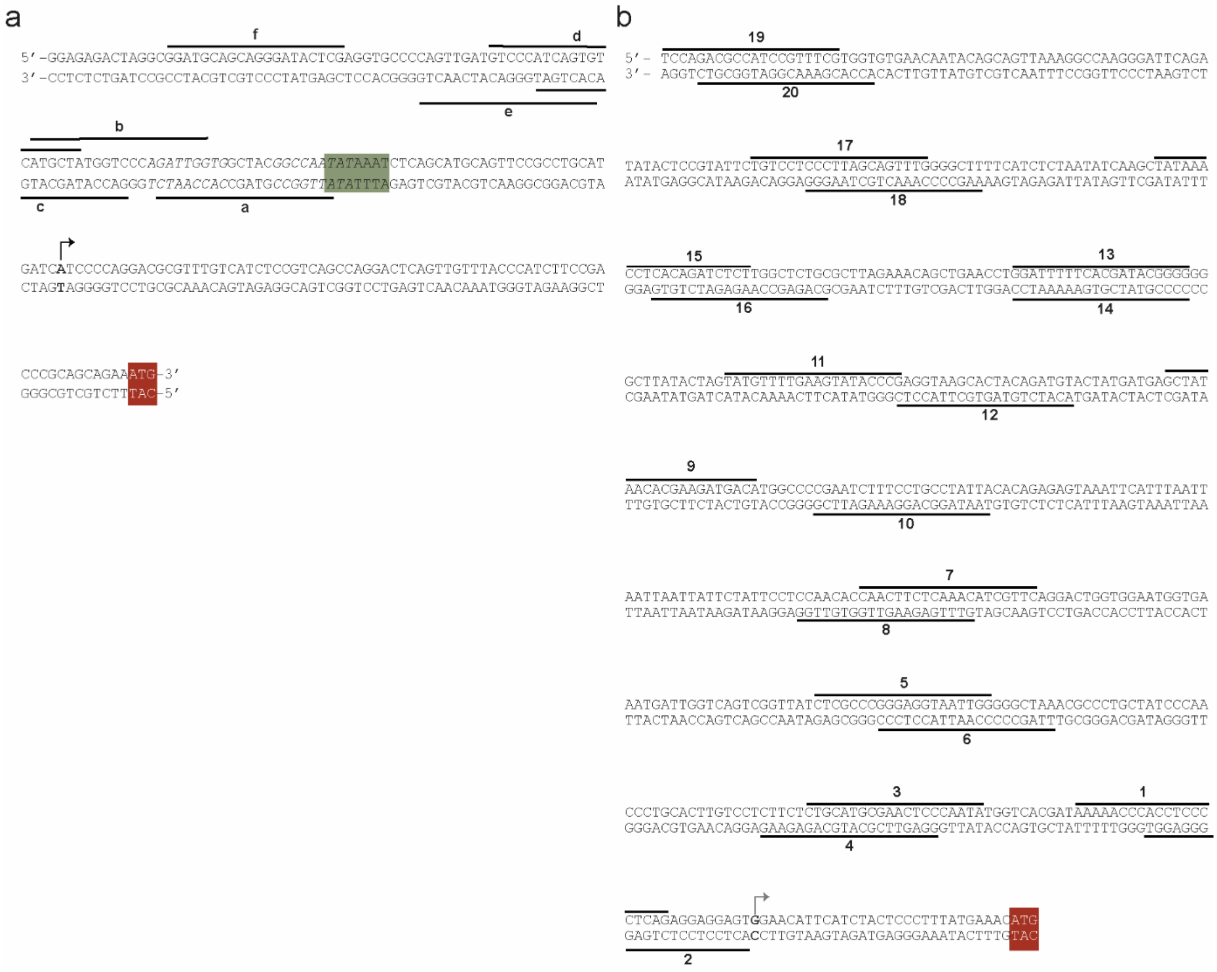

**Figure S2:** Representation of sgRNA targeting sequences on promoter sequences of *penDE*-CP **(a)** and *macR* **(b)**. Transcription start site (TSS) of *penDE* indicated as black arrow, predicted TSS of *macR* indicated as gray arrow. Red box indicates translation start codon, green box indicates TATA-box.

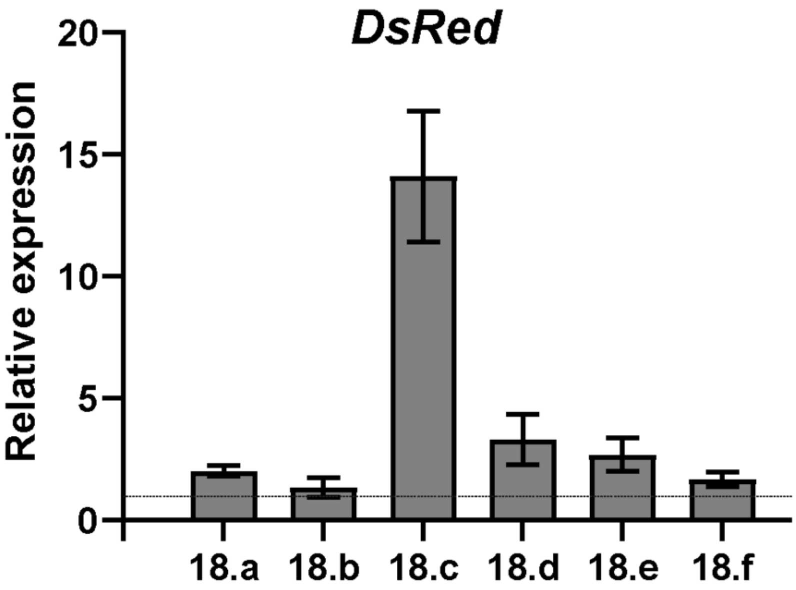


**Figure S3:** qPCR analysis showing expression levels of *DsRed* in CRISPRa strains relative to strain carrying pAMA18.0 no-sgRNA negative control (dotted line) after 5 days of growth in SMP.

**a)**

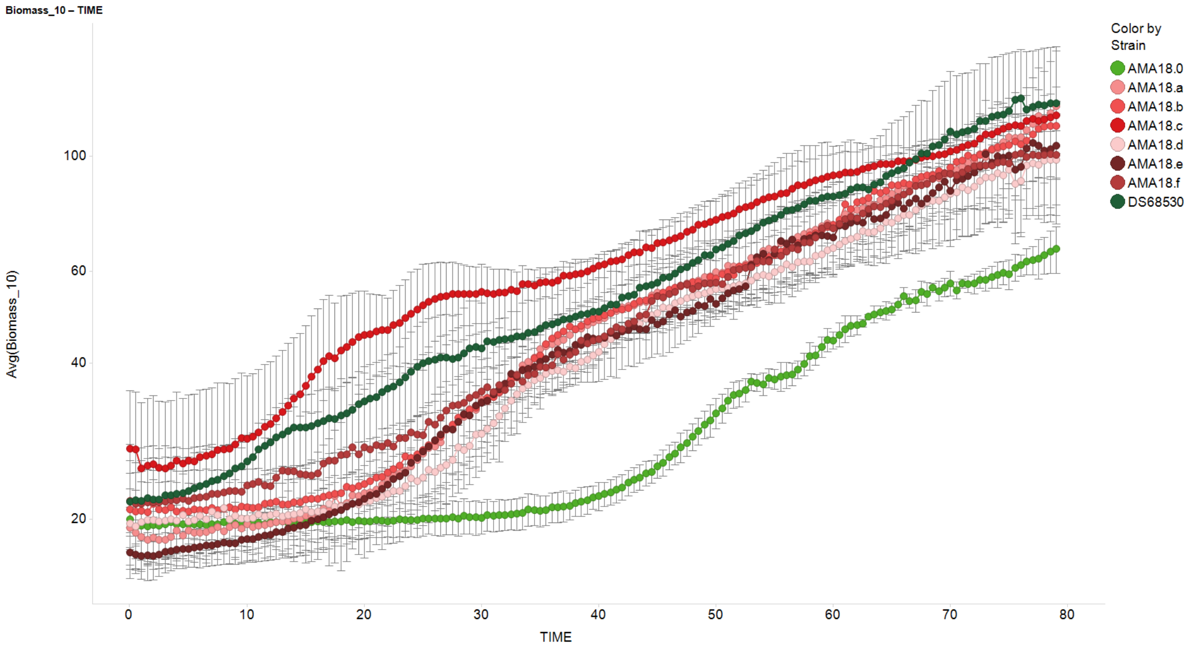
Biomass_10 – TIME
Avg(Biomass_10)
100
60
40
20
0
10
20
30
40
50
60
70
80
TIME
Color by
Strain
AMA18.0
AMA18.a
AMA18.b
AMA18.c
AMA18.d
AMA18.e
AMA18.f
DS68530

**b)**

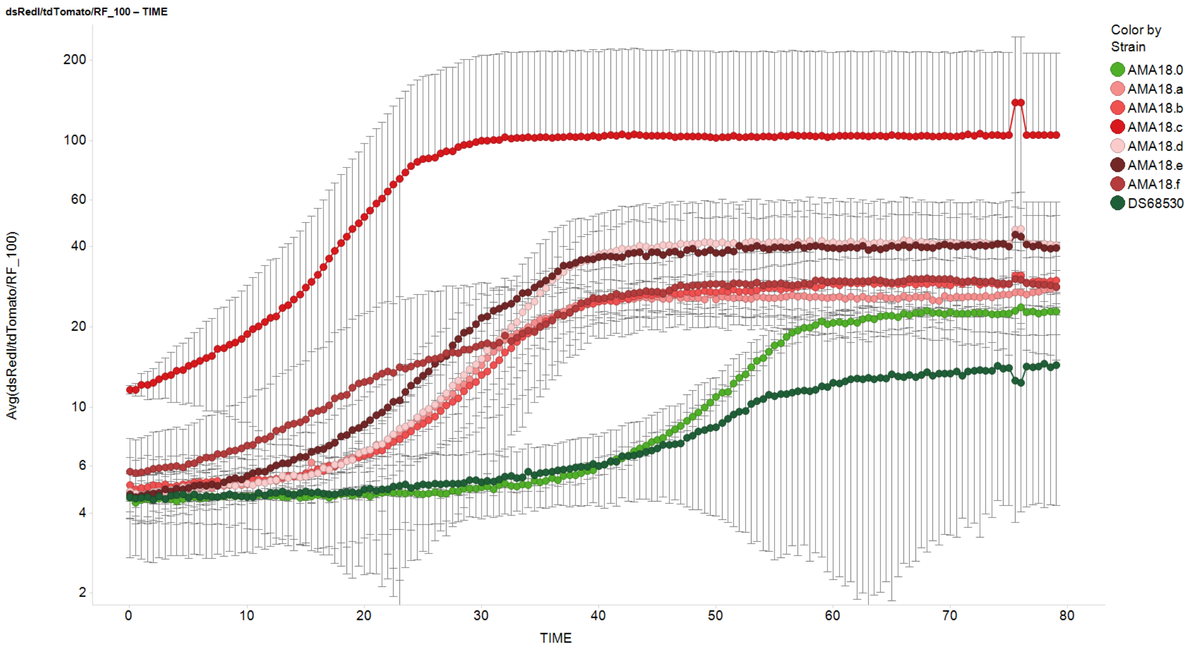
dsRedl/tdTomato/RF_100 – TIME
Avg(dsRedl/tdTomato/RF_100)
200
100
60
40
20
10
6
4
2
0
10
20
30
40
50
60
70
80
TIME
Color by
Strain
AMA18.0
AMA18.a
AMA18.b
AMA18.c
AMA18.d
AMA18.e
AMA18.f
DS68530

**c)**

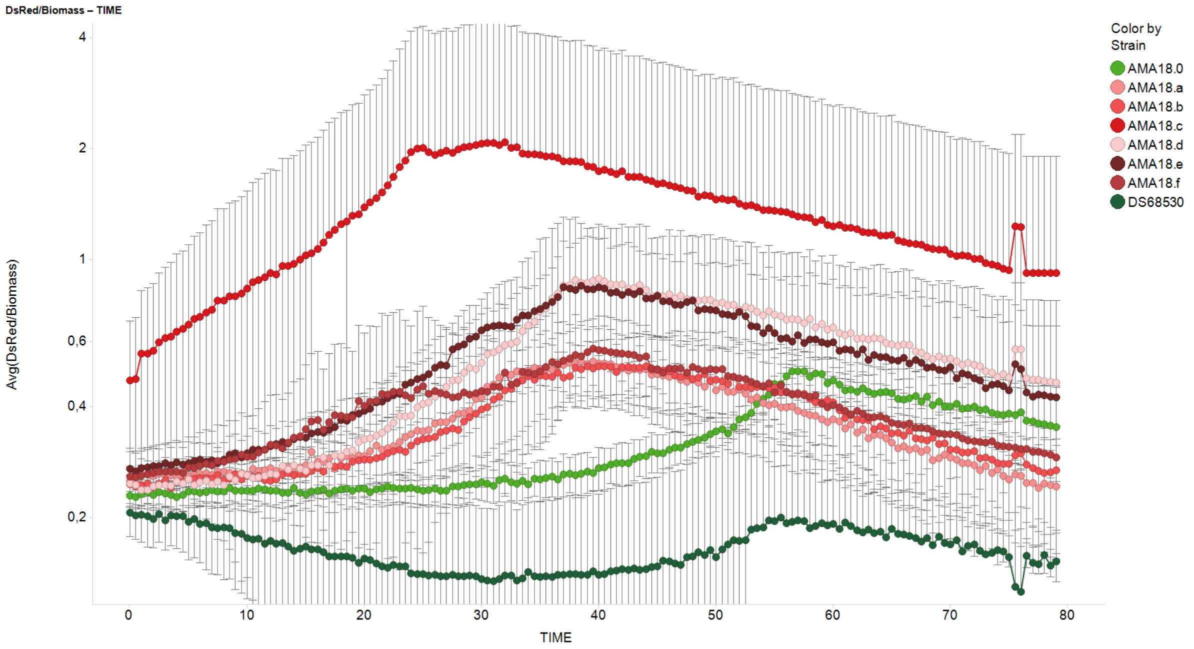


**Figure S4:** Development of **(a)** biomass **(b)** DsRed fluorescence **(c)** and DsRed fluorescence corrected for biomass for indicated CRISPRa strains in BioLector microbioreactor, compared to non-sgRNA control (AMA18.0) and *DsRed*-free parent strain (DS68530). Strains were cultivated in SMP liquid media, supplemented with terbinafine (except parent strain DS68530). Data were obtained from 3 separate experiments, each consisting of 2-3 biological replicates; error bars show the standard deviation.

**> *macR*_(*Pc16g00410*)_ *Penicillium_rubens*_DS68530_complete_cds**

atgcctttatccgagccccgccaaatcaagaaaccccggttatcgctcagttgcatcgtctgccgacgacgcaaggttcgctgcggtcgcgaacagccagaatgtgccaactgtgtgcgcatgaa agagaattgtgtctaccgggccatggtccgtgac
gagtccaccggccgtgtacgaccagtatcaccgcaggacaaagactccagggactctacatcggatgcccgtccagacctctcttggccgcattggg gccgagacaccaatgccccataccccgataagccgtctaggcctccttctcgtcctcaagta
tccgccaatgccagtccatcgccgcaacgacaagagccatacccaacagtcccttcatgggaagaggctatacaactcccaagttatcgcgatgcgaatgctcctcaaattaatggtagctcgac aactacccgcgatccctcccctgcgccatcaaca
agattattccgggccccgagtgagcctccctgccgcgattatttgagtatacgacgaggtggtcgcgtgcgatacgttggtcagactttctgggggttcgttgctgggaaggaaagtctgagcgatgatttcttcgacgaaaaccgtcatgcccaccccg at
ctccccttccacatatttcctcgatggggatgttcaatctcctgcggtcgctgccgaccaaacccgttagtgataccctgctcgagactttttccttgccgtatggccacttgttcccctcct acatccgccttccctgcaggcagactacgacgagttttggg
aatggtgtcggaatagcgagaattctttaccttcggataaactccgcgatgacccaaccttgatctgcctgctctttgcagtcctatactgtggcgcatctgccgcgccggcagccagctgggtg aacaccaacctgcagggtctacagaaggagacgac
agtgagccatctcaagtcggcatatacaacgagtctttctctatgtcaatatcaggaac atccaactttaaatacgctggtttcgaatttgctgacgggaccgtttctggatcggccgttcgagcccatgcgtagcctggttaacgtgagcaccacggtgcgca
ttgcccagactatggggttgcatcgggagggggtaggatctgcgctgagttccgttgatcgggaaatgcggcgcccgggtctggtggcacattatttggcttgatgtgcagtcgagcatctccacg gggctaaccccctgttgcgggaacgaggccttgga
tgcggtgggcatggttggcactgacaatgcagagccgagcgatatccccgctggaatttccccacccaacgaactggtgactaacagacagtcggtagccatgctatatgctatcgggcgcttcc aggctgctcgcttacaggcgaggaccgtggcgc
acctgcagagtgcgcatggtccaagccagcatggatttggcgagctgatcacagatgccaaggagcttctgcagaagattgactcgctt atcgcacgcgttccaacacaggggatccccgaaatgggatacataccatcccgcctggcgaatgcgtctc
cgtccacccagcccttgttgtacaaggacgattcaagccagccaaccgtctttgcagcatggacgcgaatcatgctaacattgctgaaatcagaaatggctatcttattgcagaaaccatttctt ccacccccggacagtgcgaacccgcaatcgcgcaag
tcatggaccagtatggcgcagctctgcgtgaattacttgcgtatttacctgcagctgtatcaggcccctgctttctccccatacgcgtggttctgctgcagccactatgggcctctccaatgcgtattcatcaccctaatgtacctccat tacttccagcactctgg
agagaccacgctggcccgatattgtgttgatgaggttatacatcactgcgtcgcccagtatcaagctccaggaccttcctcgacaaggactagccttgatggtactgattccag tgggggcaaaatgccaatgccattggccattcaggttctcgttgacctt
cacgaacggctcgactcgtctctcggacctgaagacaaggctccgccactggacctaatcgagtgtcaggcccgattttctatgtctcaccttgctaccaaggcgtccaacttgcgcgcgacgtc tgaccagccttcttctgatgtatcgtctaccactcgca
cccaccacaattgcgagacgcctcccataaccaccggggctcctccagttgctgctggtaacaaatcaatcccaccgaacagcgtctttgtggccgggagtgactcaggcttggatatggatttccttgctacgatttcggatcttgaggcctggtcctcgtc
attgattctggaatccgacaatctcctcgcacgtcctgataatatgacgcctgatcatgctgtaatcacagggttaggctcacaaagtaccgcgactggtcgccgtggccttcccggcggactcg attttccagtgtaa

**Figure S5:** Complementary DNA sequence of *macR* (*Pc16g00410*) of *Penicillium rubens* DS68530 as determined by Sanger sequencing.

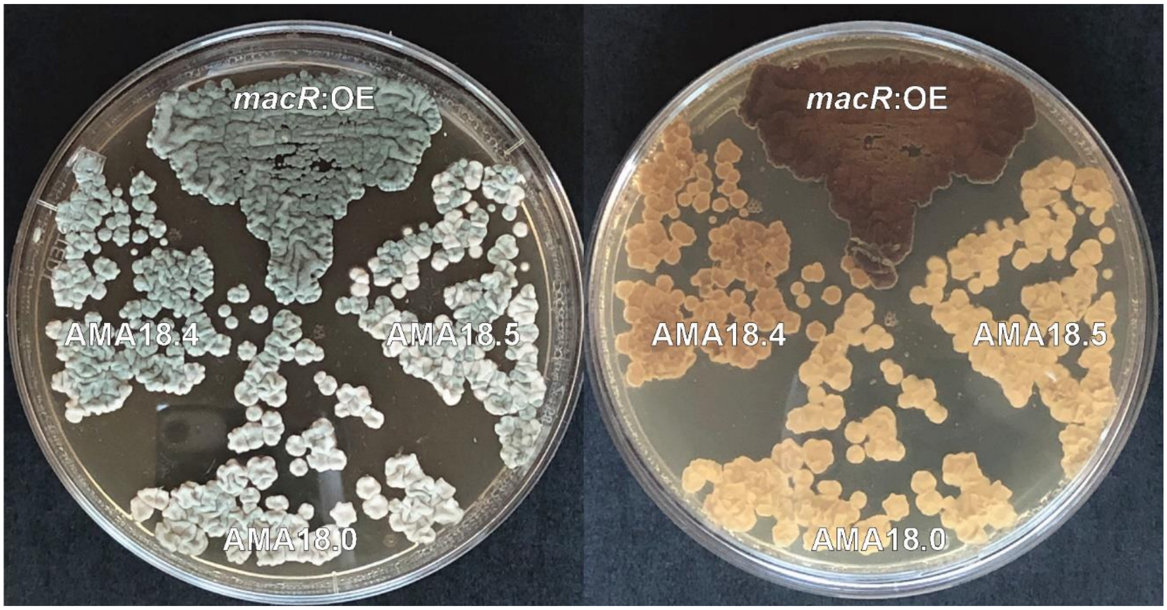


**Figure S6:** Dark pigmentation of hyphae due to *macR* overexpression in *Penicillium rubens* DS68530 *macR:OE* and in the CRISPRa AMA18.4 strain after 5 days of cultivation on R-agar.

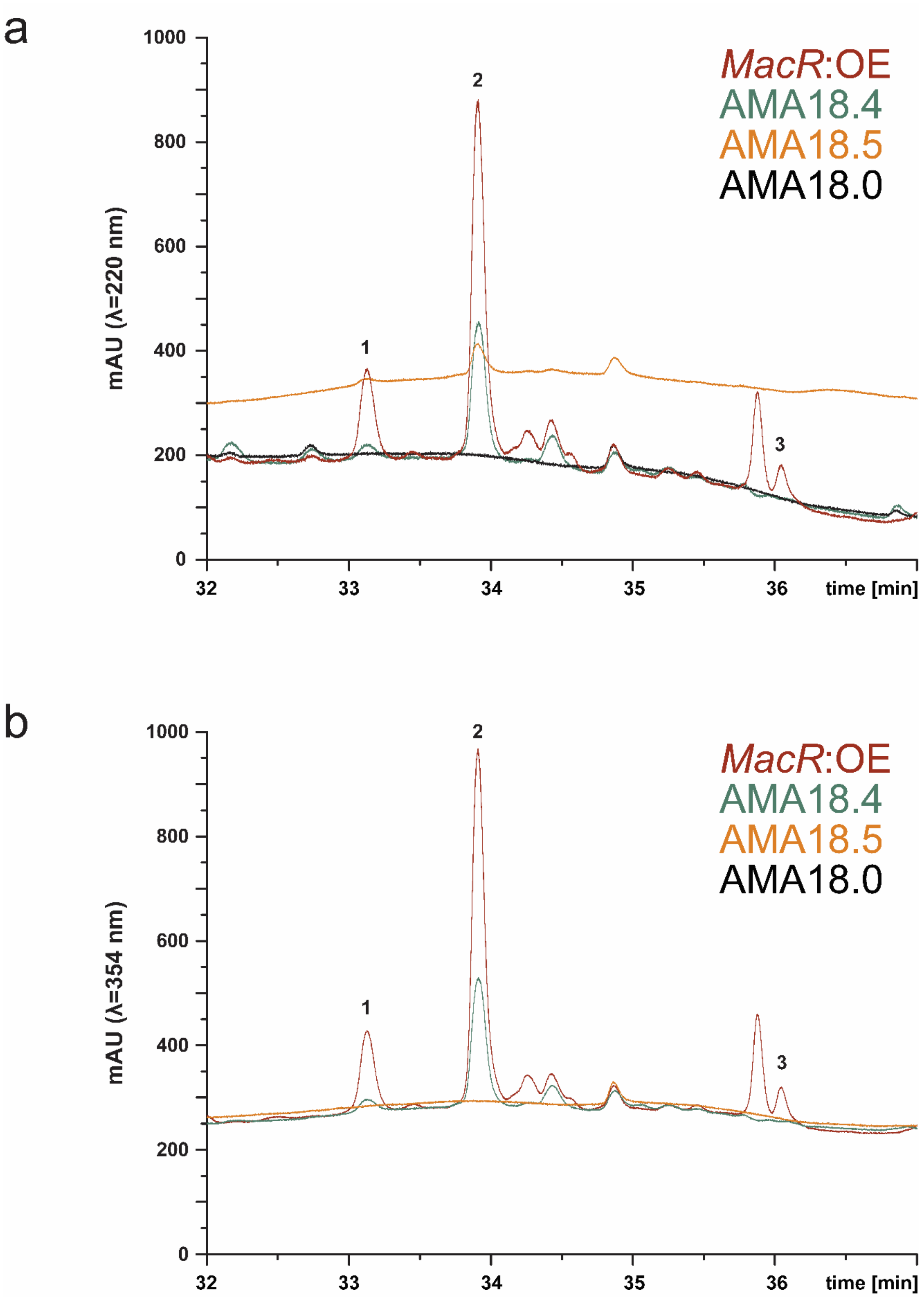


**Figure S7:** LC-MS UV-VIS chromatograms of hyphae extracts of CRISPRa and *macR*:OE strains analyzed at (**a)** λ=220 nm and (**b)** λ=354 nm.
